# NEDDylation stabilizes eIF3g and eIF3i during stress

**DOI:** 10.64898/2026.08.22.746433

**Authors:** Aravinth Kumar Jayabalan, Ramesh Mariappan, Abirami Rajendiran, Takbum Ohn

## Abstract

Stress granules (SGs) are cytoplasmic biomolecular condensates that assemble when translation initiation stalls, sequestering stalled preinitiation complexes and associated RNA-binding proteins. How individual initiation factors are targeted to SGs and released following stress recovery to reinitiate translation remains poorly understood. Here, combining a NEDD8-conjugate proteome with our previously reported arsenite-induced NEDD8 interactome and curated RNA granule databases, we find that eIF3g and eIF3i are shared, high-confidence NEDDylated SG components. NEDDylation of eIF3g and eIF3i-associated complexes is readily detected at steady state and declines under arsenite stress. Intriguingly, only full-length eIF3g is recruited to SGs. eIF3g lacking the RRM domain strongly inhibits SG formation, whereas the RRM domain alone neither inhibits SG assembly nor localizes to SGs. Blocking the NEDD8 pathway— by NAE inhibition with MLN4924, depletion of NEDD8 pathway components, or expression of the deNEDDylase NEDP1—accelerates the loss of eIF3g and eIF3i protein during stress. Our data indicate that NEDDylation marks a degradation-resistant pool of eIF3g/eIF3i that is competent for SG localization, linking the NEDD8 pathway to initiation-factor proteostasis and condensate partitioning, and potentially making these factors available for translation reinitiation during stress recovery.

## Introduction

Cells respond to environmental stress by rapidly reprogramming translation^1^. Phosphorylation of the α-subunit of eIF2 (eIF2α) limits ternary-complex recycling, stalls 48S preinitiation-complex formation, and drives the assembly of SGs—non-membranous biomolecular condensates enriched in mRNA and initiation factors^2^. The RNA-binding proteins G3BP1 and G3BP2 nucleate the core protein-protein and protein-mRNA interaction network, and the duration, intensity, and type of stress determine which factors are recruited to SGs^3,4^; arsenite and sodium chloride are widely used to induce canonical and non-canonical SGs, respectively^5^. Although SG composition has been extensively catalogued^6–8^, the mechanisms that select specific factors for partitioning and for translation reinitiation once the stress is resolved remain poorly defined. Dysregulated assembly and disassembly of SGs are prevalent in age-related diseases, including cancer and neurodegeneration^9–11^.

Post-translational modification provides much of this selectivity. O-GlcNAcylation of ribosomal proteins is required for SG aggregation but not for translational arrest^12^; ADP-ribosylation regulates SG assembly in the cytoplasm^13,14^, with PARP10 acting on G3BP1 to initiate condensation and translation arrest^15^ and removal of the mark by a viral ADP-ribosylhydrolase driving SG disassembly^16^. SUMOylation and site-specific ubiquitination of G3BP1 likewise promote SG disassembly in a context-dependent manner^17,18^. Other post-translational modifications are known to regulate SG dynamics. These modifications act at separable steps, including translational shutoff, condensation, and dissolution, indicating that SG dynamics are governed not by a single switch but by a set of marks on distinct substrates^19^.

NEDD8 is a ubiquitin-like protein whose best-characterized role is activation of cullin-RING ligases, but a growing number of non-cullin substrates have been described^20^, and NEDDylation can alter substrate stability, localization, and interactions^21^. Ribosomal proteins were among the first non-cullin NEDD8 targets identified, where the modification opposes their degradation, placing the pathway at the translational apparatus itself^22,23^. We previously reported that the NEDD8 pathway is required for SG assembly under oxidative stress and identified the splicing factor SRSF3 as one of the NEDDylated SG regulators^24^. More recently, the NEDD8 pathway has been linked to the opposite arm of the SG life cycle: loss of the deNEDDylase NEDP1 accelerates SG clearance and prevents pathological solidification through hyper-NEDDylation of PARP1^25^. NEDDylation thus appears to act at multiple nodes of SG regulation, with the outcome determined by substrate-level specification.

Here, we intersect a proteome-wide map of endogenous NEDD8 sites obtained from deconjugase-deficient cells^26^, a curated RNA-granule database^27^, and our arsenite-induced NEDD8 interactome^24^, converging on the eIF3 subunits eIF3g and eIF3i. Both are peripheral to the functional core of mammalian eIF3 and dispensable for reconstituted initiation complex assembly, which makes them plausible candidates for regulated partitioning^28–30^. We show that eIF3g is a canonical NEDD8 substrate whose modification is most abundant at steady state and declines during arsenite stress. Disrupting the pathway—by NAE inhibition, deNEDDylase expression, or depletion of NEDD8 pathway components— accelerates loss of both eIF3g and eIF3i protein levels, most markedly under stress. Domain analysis indicates that eIF3g NEDDylation occurs in the lysine-rich N-terminal region, as previously reported^26^. Intriguingly, eIF3g localization to SGs is determined by the RRM domain, whose deletion strongly inhibits SG formation; however, the RRM domain alone neither localizes to SGs nor affects SG formation. We propose that NEDDylation maintains a degradation-resistant pool of eIF3g and eIF3i that remains competent for condensate recruitment, linking the NEDD8 pathway to initiation-factor proteostasis and translation reinitiation following stress recovery.

## Results

### eIF3g and eIF3i are high-confidence NEDDylated SG proteins

We previously reported the arsenite-induced NEDD8 proteome dataset and showed that the NEDDylation pathway plays a key role in SG assembly by characterizing SRSF3, a splicing factor, as one of its substrates^24^. Importantly, our dataset also contained several ribosomal proteins, consistent with a previous report that identified these proteins at steady state^22^. This prompted us to identify specific SG proteins that are NEDDylated at steady state. To determine these proteins, we intersected our previously reported NEDD8-modified SG-associated protein dataset^24^ with a NEDD8 proteome dataset^26^ and identified 66 proteins that were shared between these two datasets, representing candidates that are

NEDDylated at steady state (**Fig. 1A, Supplementary Table S1**). Functional classification of this overlap showed a strong enrichment for translation-related factors, with 40S and 60S ribosomal subunits together constituting the largest categories, followed by metabolic, RNA-binding, and cytoskeletal proteins, translation factors, and ubiquitin–proteasome/NEDD8 components (**Fig. 1B**).

**Figure 1.**
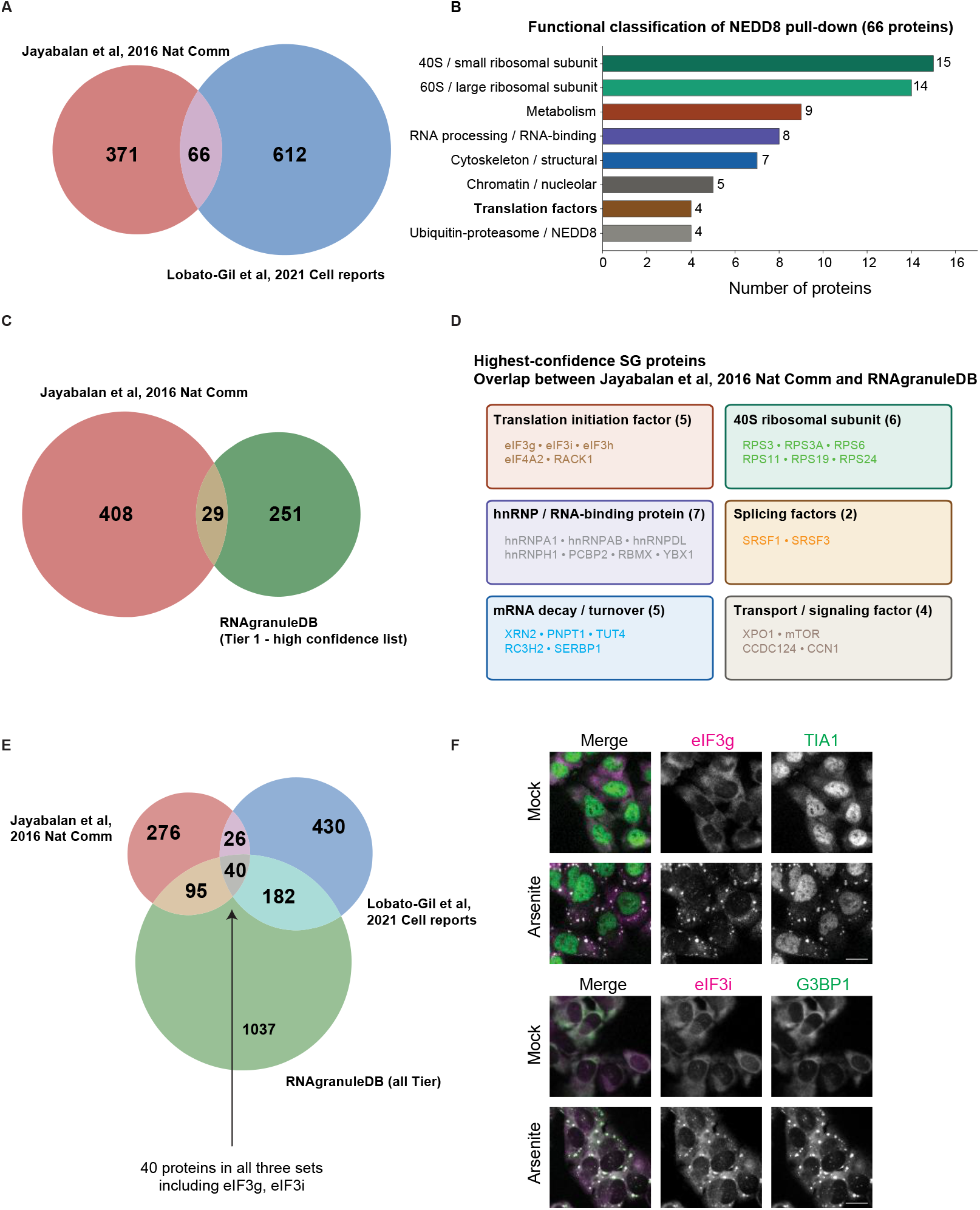
eIF3g and eIF3i are high-confidence SG components and candidate NEDD8 substrates. (**A**) Venn diagram of the overlap between the NEDD8-modified SG-associated protein dataset (Jayabalan et al., 2016) and a NEDD8 proteome (Lobato-Gil et al., 2021). (**B**) Functional classification of the 66 shared proteins (number of proteins per category indicated). (**C**) Overlap between the NEDD8-modified SG-associated protein dataset (n=437) and the RNAgranuleDB Tier 1 (high-confidence, n=280) list, yielding 29 proteins. (**D**) The 29 shared proteins from (C) grouped by function. (**E**) Three-way overlap between the NEDD8-modified SG interactome, the NEDD8 proteome, and all tiers of RNAgranuleDB; 40 proteins are common to all three sets, including eIF3g and eIF3i. (**F**) Representative immunofluorescence of eIF3g (magenta) with TIA1 (green) and eIF3i (magenta) with G3BP1 (green) in U2OS cells, mock-treated or treated with 0.5 mM arsenite for 1 h. Scale bar, 10 µm.

Next, to restrict this list to the high-confidence SG components, we intersected our NEDD8-modified SG-associated protein dataset with the Tier 1 (high-confidence) list of RNAgranuleDB, yielding 29 shared proteins (**Fig. 1C, Supplementary Table S1**). These included translation initiation factors (eIF3g, eIF3i, eIF3h, eIF4A2, RACK1), 40S ribosomal subunits, hnRNP/RNA-binding proteins, splicing factors (including SRSF3), mRNA-decay factors, and transport/signaling factors (**Fig. 1D**). A three-way intersection of the NEDD8-modified SG interactome, the NEDD8 proteome, and all tiers of RNAgranuleDB identified 40 proteins common to all three datasets, including eIF3g and eIF3i (**Fig. 1E, Supplementary Table S1**).

We focused on eIF3g and eIF3i because these eIF3 subunits, present in all three datasets, are dispensable for the assembly of the translation initiation complex^28,29^. Consistent with their annotation as SG proteins and as hits from a previously identified SG proteome^6^, endogenous eIF3g and eIF3i readily redistributed into cytoplasmic puncta that colocalized with the SG markers TIA1 and G3BP1, respectively, upon arsenite treatment, whereas they showed diffuse cytoplasmic staining in mock-treated cells (**Fig. 1F**). We further confirmed the SG localization of several other initiation factors under arsenite, thapsigargin, and clotrimazole stress (**Fig. S1**). Together, these data identify eIF3g and eIF3i as candidate NEDD8 substrates and bona fide SG components.

### eIF3g is NEDDylated at steady state and the modification declines during stress

To test directly whether eIF3g and eIF3i are NEDDylated, we co-expressed FLAG-tagged eIF3g with 6XHis-tagged NEDD8 or a conjugation-defective 6XHis-NEDD8ΔΔ mutant and captured NEDD8 conjugates by His pull-down under denaturing conditions. A NEDDylated FLAG-eIF3g species was observed in wild-type NEDD8-transfected cells but not in vector- or NEDD8ΔΔ-transfected cells, confirming eIF3g as a canonical NEDD8 substrate (**Fig. 2A**). Surprisingly, we observed a decrease in the eIF3g NEDDylation mark upon arsenite treatment (lane 2 vs. lane 5). Because both our proteome^24^ and the dataset of Lobato-Gil et al^26^ contained additional translation factors, we asked whether this reduction in NEDDylation during arsenite stress was eIF3g-specific or extended to translation complexes more broadly. To this end, we performed a reciprocal experiment in which we immunoprecipitated FLAG-eIF3i under non-denaturing conditions to isolate intact translation complexes, and blotted for NEDD8. We observed a similar trend to that for eIF3g, in which NEDDylation of eIF3i and its associated complexes was reduced under stress (**Fig. 2B**). In both experiments, the modified species were most abundant in mock-treated cells and decreased following arsenite treatment, suggesting deNEDDylation of these factors under arsenite stress (**Fig. 2A, B**).

**Figure 2.**
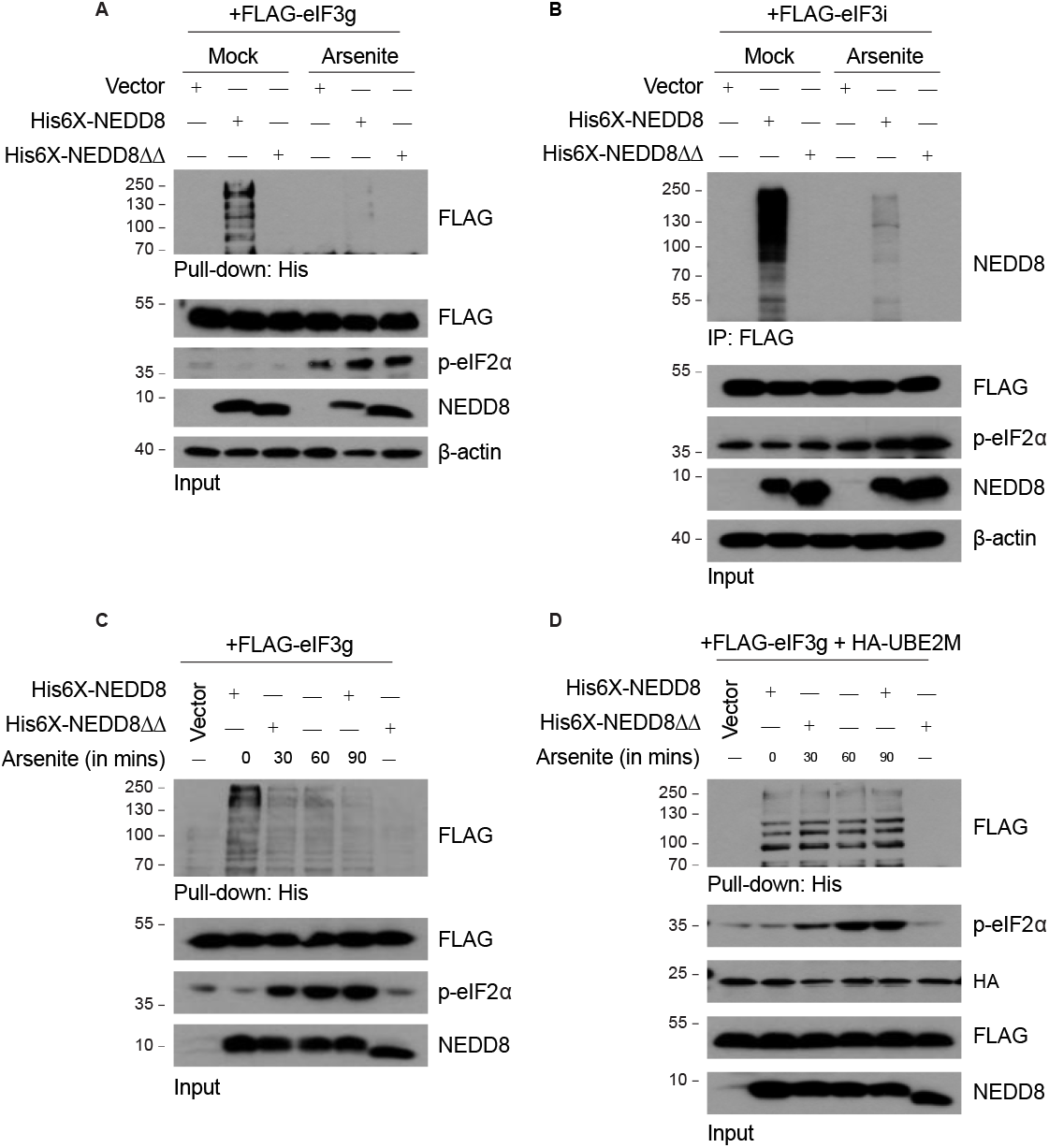
eIF3g is a novel NEDDylation substrate. (**A**) His pull-down of NEDD8 conjugates from 293T cells co-expressing FLAG-eIF3g with vector, 6XHis-NEDD8, or conjugation-defective 6XHis-NEDD8ΔΔ, under mock- and arsenite-treated conditions (0.5 mM for 60 min). Pull-down samples were probed with anti-FLAG antibody. (**B**) FLAG immunoprecipitation under non-denaturing conditions from 293T cells expressing FLAG-eIF3i probed for NEDD8, under mock- and arsenite-treated conditions (0.5 mM for 60 min). (**C**) His pull-down of NEDD8 conjugates from FLAG-eIF3g-expressing 293T cells across an arsenite time course (0–90 min). (**D**) His pull-down across an arsenite time course (0–90 min) from 293T cells co-expressing FLAG-eIF3g and HA-UBE2M.

Next, we performed a time course experiment with arsenite treatment and observed that NEDDylated eIF3g was maximal in untreated cells and declined progressively over 30–90 min of arsenite exposure, confirming a time-dependent phenomenon (**Fig. 2C**). This decrease in NEDDylation was rescued by co-expression of the NEDD8-conjugating E2 enzyme HA-UBE2M, which sustained eIF3g NEDDylation across the arsenite time course. This is consistent with the modification proceeding through the canonical NEDD8 E2 enzyme^20^ and suggests that higher expression of the E2 conjugating enzyme maintains the NEDDylated eIF3g pool at basal level under stress (**Fig. 2D**). Together, these data establish that eIF3g is a bona fide NEDD8 substrate whose modification is predominant in unstressed, steady-state conditions and declines during arsenite stress.

### eIF3g NEDDylation maps outside the RRM, whereas the RRM directs SG partitioning

eIF3g contains 29 lysines distributed across an N-terminal eIF3i/eIF4B-binding region (21 lysines), a CCHC zinc-finger (four lysines: K150, K153, K161, K172), and a C-terminal RRM domain (four lysines: K272, K274, K280, K316) (**Fig. 3A**). To localize the modification and the domain responsible for SG localization, we generated FLAG-tagged eIF3g truncations: full-length (WT), a deletion lacking the RRM domain (ΔRRM), and the RRM domain alone (RRM only) (**Fig. 3B**). Previous work mapped three NEDDylation sites in eIF3g— K193, K195, and K209—all of which lie N-terminal to the RRM domain, in the region between the CCHC zinc finger and the RRM^26^. In His-NEDD8 pull-downs, we observed that both eIF3g WT and ΔRRM were heavily NEDDylated, consistent with the number of lysines retained in these constructs. In contrast, the RRM-only construct showed negligible modification, in line with the reported eIF3g modification sites (**Fig. 3C**).

**Figure 3.**
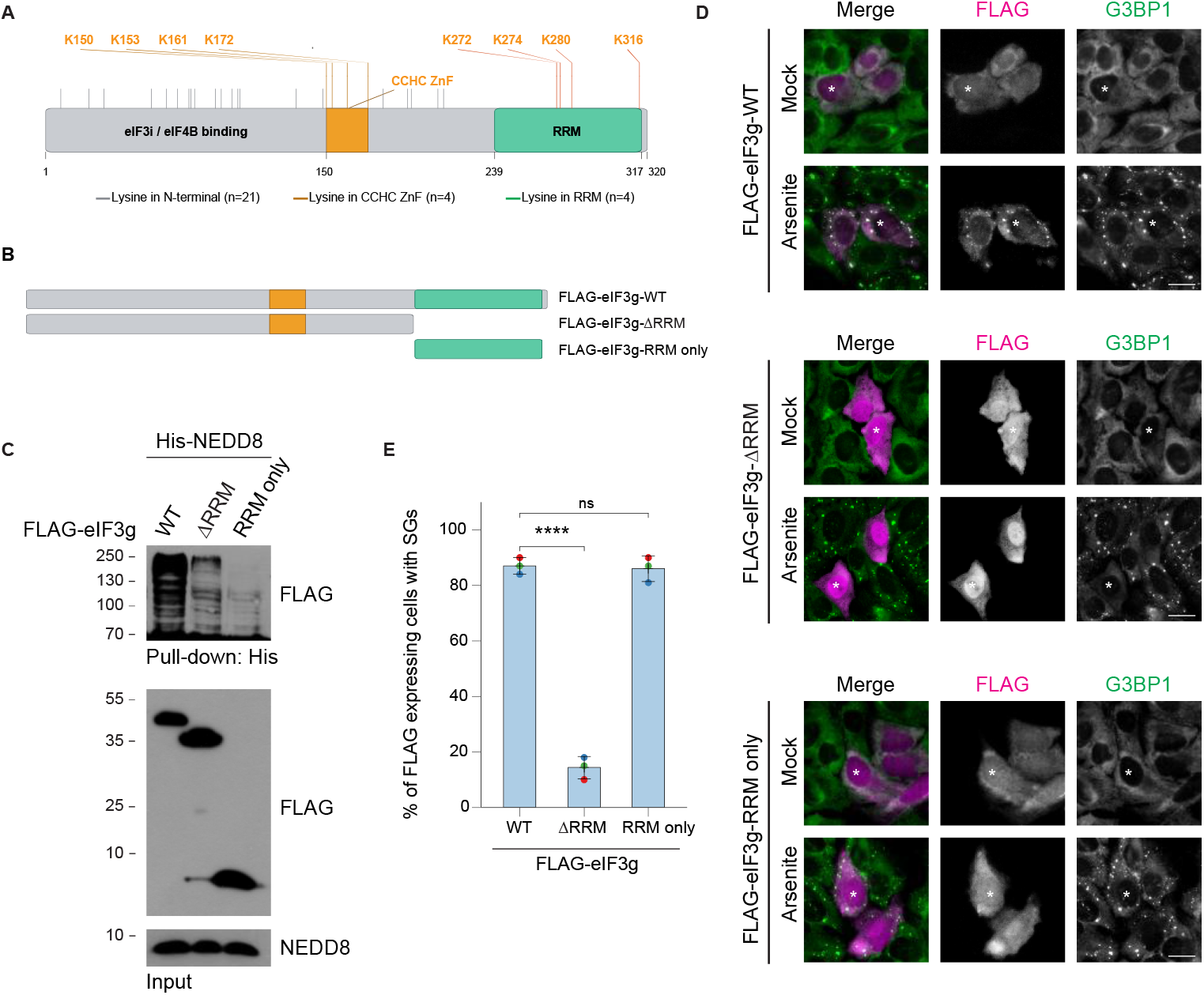
The eIF3g RRM domain is required for SG recruitment. (**A**) Domain architecture of eIF3g with lysine positions indicated. (**B**) Schematic of FLAG-eIF3g constructs. (**C**) His-NEDD8 pull-down of FLAG-eIF3g WT, ΔRRM, and RRM-only constructs expressed in 293T cells, probed with an anti-FLAG antibody. (**D**) Representative immunofluorescence of FLAG-eIF3g WT, ΔRRM, and RRM-only (magenta) with G3BP1 (green) in mock- and arsenite-treated cells at 0.2 mM for 60 min. Scale bar, 10 µm. (**E**) Percentage of FLAG-expressing cells containing G3BP1-positive SGs. ****P < 0.001, two-tailed, unpaired Student’s *t*-test.

We next asked whether these constructs display differential recruitment into SGs, given that the RRM-only construct lacks a mapped modification site. Intriguingly, only eIF3g-WT was efficiently recruited to G3BP1-positive SGs upon arsenite treatment, whereas eIF3g-ΔRRM strongly inhibited SG formation and remained largely diffuse in the cytoplasm (**Fig. 3D**). In eIF3g-RRM-only-expressing cells, the construct neither localized to SGs nor inhibited their formation. Quantification showed that ~85% of eIF3g-WT and RRM-only–expressing cells contained G3BP1-positive SGs, compared with ~15% of ΔRRM-expressing cells (**Fig. 3E**). Altogether, these data indicate that the mapped NEDDylation sites lie outside the RRM and that the RRM is required for eIF3g partitioning into SGs.

### NEDDylation protects eIF3g and eIF3i from degradation

NEDDylation of ribosomal proteins helps to stabilize them under stress conditions^22^. The reduction in NEDDylation of eIF3g and of associated translation complexes under stress prompted us to hypothesize that the modification might play a protective role for eIF3g and eIF3i. To examine eIF3g/eIF3i protein stability, we performed cycloheximide (CHX) chase experiments to determine the degradation rates of eIF3g and eIF3i^31^. First, we pretreated cells with either DMSO or the NAE inhibitor MLN4924^32^ and then chased with CHX alone or with CHX plus arsenite. Our western blot analysis revealed that eIF3g and eIF3i declined more rapidly when NEDDylation was blocked with MLN4924, an effect that was heightened by concurrent arsenite treatment (**Fig. 4A, B**).

**Figure 4.**
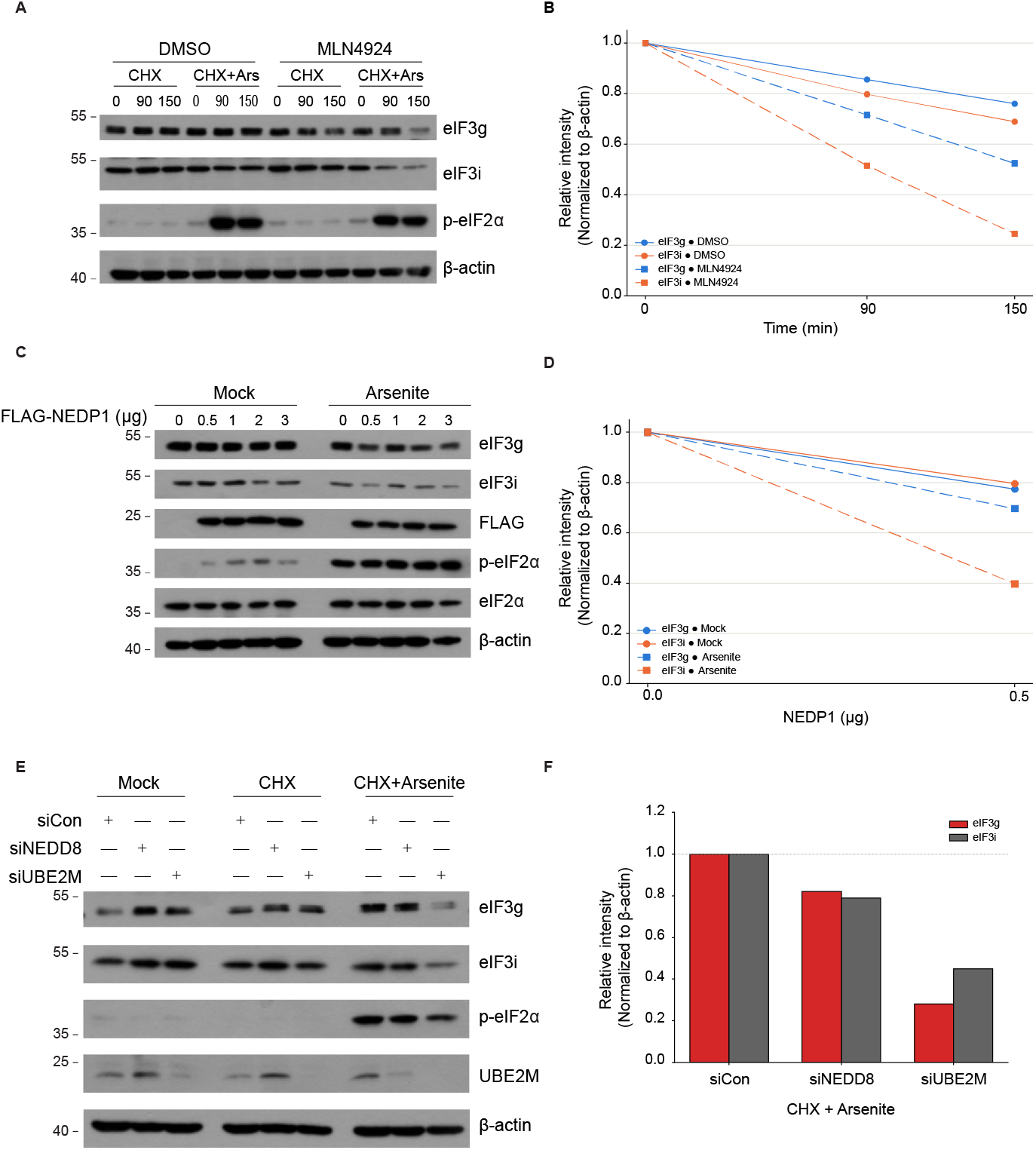
NEDDylation protects eIF3g and eIF3i from degradation. (**A**) Immunoblot of eIF3g and eIF3i from U2OS cells treated with DMSO or MLN4924 and subjected to cycloheximide (CHX) chase without or with 0.2 mM arsenite (CHX+Ars) for 0, 90, and 150 min. (**B**) Quantification of eIF3g and eIF3i intensity from (A) normalized to β-actin. (**C**) Immunoblot of eIF3g and eIF3i in mock- and 0.2 mM arsenite-treated U2OS cells expressing increasing amounts of FLAG-NEDP1 (0–3 µg). (**D**) Quantification of eIF3g and eIF3i intensity from (C) normalized to β-actin. (**E**) Immunoblot of eIF3g and eIF3i in U2OS cells transfected with control, NEDD8, or UBE2M siRNA under mock, CHX, and CHX + 0.2 mM arsenite conditions. (**F**) Quantification of eIF3g and eIF3i intensity from (E) normalized to β-actin.

Consistent with a stabilizing role for the modification, dose-dependent expression of the deNEDDylase FLAG-NEDP1 reduced steady-state levels of eIF3g and eIF3i, with the reduction more pronounced under arsenite than in mock-treated cells (**Fig. 4C, D**). Finally, depletion of NEDD8 or UBE2M by siRNA lowered eIF3g and eIF3i levels relative to control siRNA, most evidently under CHX and arsenite co-treatment conditions (**Fig. 4E, F**). These data suggest that NEDDylation marks a degradation-resistant pool of these initiation factors, and that disrupting NEDDylation by NAE inhibition, deNEDDylase overexpression, or depletion of pathway components results in the degradation of eIF3g and eIF3i during stress.

### NEDD8 pathway activity is required to mobilize eIF3g and eIF3i into SGs

Having established that the NEDDylated pool represents the degradation-resistant species (**Fig. 4**), we asked whether the NEDD8 conjugation pathway is required for recruitment of eIF3g and eIF3i to SGs. In DMSO-treated cells, both were recruited to arsenite-induced cytoplasmic puncta, whereas inhibition of the NEDD8-activating enzyme with MLN4924 markedly reduced SG formation, and the residual granules recruited little eIF3g and eIF3i (**Fig. 5A**). Polysome disassembly is a prerequisite for SG assembly, and blocking the NEDDylation pathway slows polysome disassembly, so ribosomes remain engaged in translation^24^. Consistent with impaired mobilization, disrupting the pathway by co-depletion of NEDD8 and UBE2M (siNEDD8+siUBE2M) retained both factors in heavier polysome fractions under arsenite treatment relative to control siRNA (**Fig. 5B**). This suggests that, in the absence of NEDDylation, eIF3g and eIF3i remain engaged with translating ribosomes rather than redistributing into SGs, although we cannot fully exclude an indirect contribution from slowed polysome disassembly. Finally, NEDD8 itself colocalized with G3BP1-positive SGs upon arsenite treatment and gradually dispersed during recovery at 30 and 60 min (**Fig. S2**), placing the NEDD8 system at SGs during the stress response and revealing its release as SGs resolve. Together, these data indicate that NEDD8 pathway activity is required to mobilize eIF3g and eIF3i.

**Figure 5.**
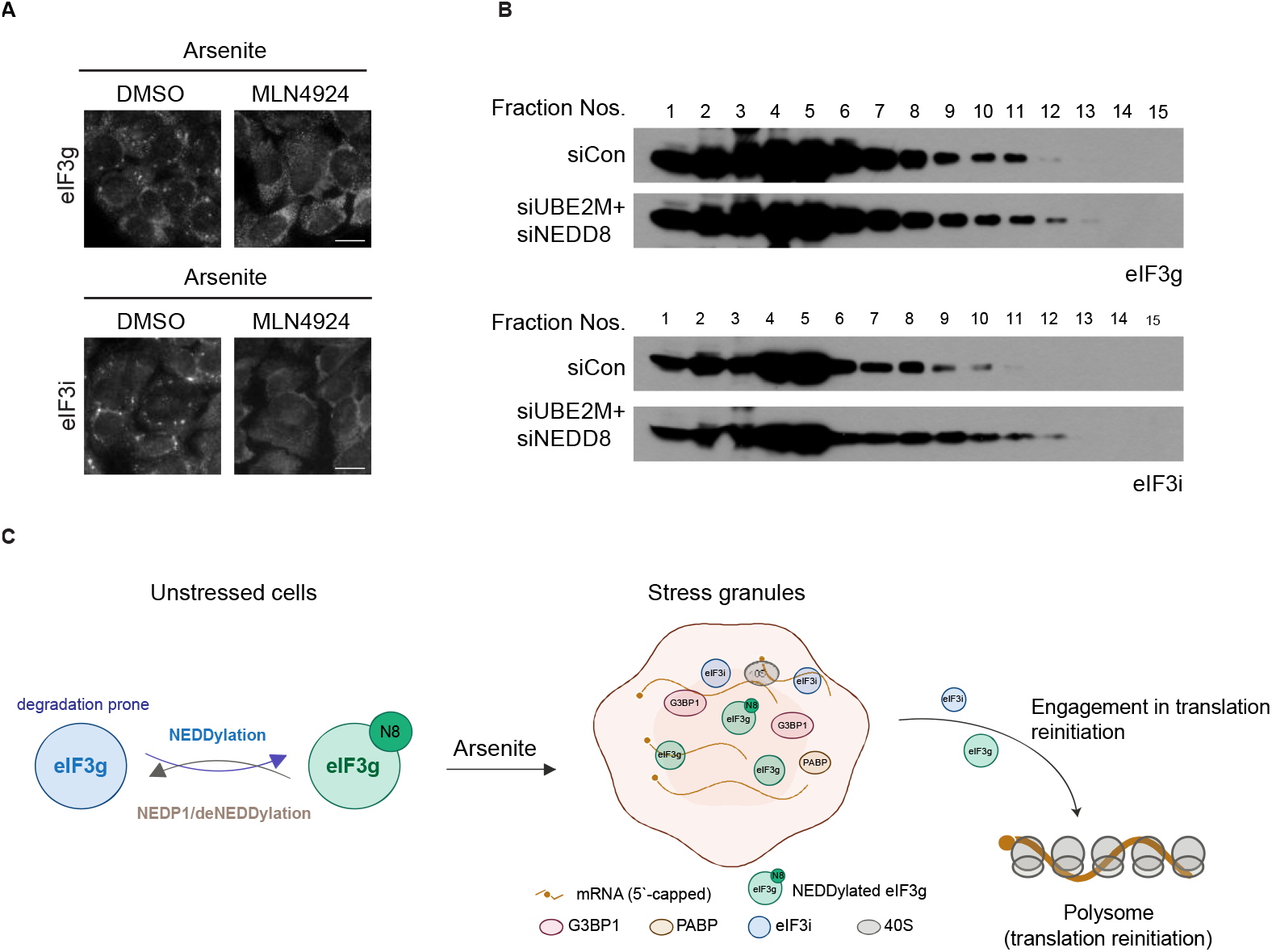
The NEDD8 pathway is required to mobilize eIF3g and eIF3i out of polysomes. (**A**) Immunofluorescence of eIF3g and eIF3i in arsenite-treated U2OS cells (0.2 mM for 60 min) pretreated with DMSO or the NAE inhibitor MLN4924. (**B**) Western blot analysis of polysome fractions from U2OS cells treated with siCon or siNEDD8+siUBE2M and subjected to arsenite treatment. A254 trace is available in Jayabalan et al, 2016. (**C**) Model: NEDDylation protects eIF3g and eIF3i under stress conditions. Scale bar, 10 µm.

## Discussion

Stress-induced reprogramming of cellular pathways is essential for cell survival, with translation control as one of the immediate responses that stall global protein synthesis^33^. Stalled translation induces transient SGs, which recruit translation factors, RNA-binding proteins, and key signaling proteins, and disassemble once stress is relieved^34^. Rapid resumption of protein synthesis after stress relief requires that the initiation machinery remain intact through the repression phase^35^. How translation factors are preserved during stress, rather than being turned over with other stalled or damaged proteins, has received little attention. Our work identifies one such mechanism: NEDDylation marks a degradation-resistant pool of eIF3g and eIF3i that is competent for SG recruitment (**Fig. 5C**).

### Translation initiation factors are novel NEDD8 substrates

We identify the eIF3 subunit eIF3g as a NEDD8-modified SG protein and implicate eIF3i and associated translation factors as likely NEDD8 substrates (**Fig. 2**). Moreover, these factors have been identified as hits in mass spectrometry analyses, and we now confirm the modification experimentally^24,26^. Mechanistically, we showed that NEDDylation protects eIF3g and eIF3i from degradation under stress conditions, and that blocking the NEDD8 pathway accelerates their loss. Intriguingly, this is the opposite of what we previously found for SRSF3^24^. We show that SRSF3 is NEDDylated only under stress and that this modification is required for efficient localization to SGs. Specifically, non-NEDDylatable SRSF3 does not localize to SGs and impairs recruitment of specific SG components, with MLN4924 treatment completely inhibiting the formation of SGs. This dual role suggests that NEDDylation serves a broader function during stress, including SG assembly and protein stabilization.

Post-translational modifications are key signaling factors that add another layer of functionality to a protein based on cell state^36^. For example, although ubiquitination is constitutively involved in signaling for protein degradation, its levels increase during cellular stress to enhance protein clearance, especially of misfolded proteins^37^. Furthermore, depending on the linkage type, ubiquitin directs a protein either to degradation or to a change in localization^38^. Similarly, our work identified roles of NEDDylation in protein stabilization and condensate assembly. While previous work identified a broader protective role for ribosomal proteins^22^, our work defines a stabilization role for translation factors that is specific to arsenite-induced oxidative stress.

### Potential link between eIF3g NEDDylation and SG recruitment

An apparent paradox is that the NEDDylated species decreases under the same stress that drives SG recruitment. One interesting question is what fraction of eIF3g is NEDDylated at steady state and in native form. Our pull-down experiment under denaturing conditions cannot directly distinguish modified from unmodified eIF3g species in cytoplasmic vs. SG fractions. Furthermore, our data suggest that eIF3g NEDDylation alone is not sufficient for SG recruitment, which is instead directed by the RRM domain (**Fig. 3**). This suggests that under stress conditions, once SGs form, a proportion of eIF3g is recruited to SGs via the RRM domain along with other NEDDylated proteins (e.g., SRSF3). These eIF3g species may be protected within SGs such that they can reinitiate translation immediately during stress recovery^39^. Testing a non-NEDDylatable eIF3g mutant would address three questions: whether NEDDylation is required for SG localization, whether translation reinitiation is normal in these cells, and the extent to which eIF3g is degraded. Because NEDD8 can form mixed chains with ubiquitin^40–42^, a non-NEDDylatable eIF3g may also block subsequent ubiquitin conjugation and proteasomal degradation.

Our findings extend an established role of the NEDD8 pathway in protein stabilization to initiation-factor proteostasis. NEDDylation has been reported to regulate the stability and localization of both cullin and non-cullin substrates, including ribosomal proteins. Work from our group and others has identified several translation factors as targets of NEDDylation, and we experimentally verified the role of eIF3g NEDDylation in protein stabilization. It would be interesting to test whether this is true for other translation factors. In addition, further research is needed to identify whether the NEDDylated translation factors are functionally competent or whether these species are inactive and are required only during stress recovery. Another approach is to test whether these factors participate in non-canonical translation, such as cap-independent translation^43,44^.

### NEDDylation during stress recovery

Several observations directly link NEDDylation to translation control. Polysome disassembly is a prerequisite for SG assembly, which is nucleated by mRNPs released from disassembling polysomes^34^. Yet, blocking the NEDDylation pathway either with the NAE inhibitor MLN4924 or by depletion of key NEDD8 pathway components slows polysome disassembly^24^. This suggests that NEDDylation of certain factors contributes to translation repression and promotes polysome disassembly. Consistent with this notion, our fractionation and western blot analysis revealed that blocking NEDDylation retains eIF3g and eIF3i in heavy polysome fractions. However, the factors that are specifically involved in translation repression upon NEDDylation remain to be identified. Furthermore, complete inhibition of SG formation requires prolonged (~24 h) MLN4924 treatment, consistent with the slower turnover of non-cullin NEDD8 substrates^45^. This suggests that non-cullin substrates are critical to drive SG assembly. Interestingly, we also observed that NEDD8 protein is released from SGs early during recovery; this may be necessary to prevent NEDDylation of other proteins within SGs.

Overall, our study identified a novel function of NEDDylation specifically during stress, where it stabilizes initiation factors; this mechanism likely regulates translation reinitiation immediately upon stress recovery.

## Materials and Methods

### Cell Culture and chemicals

U2OS (human osteosarcoma) and HEK293T cells were obtained from ATCC and maintained in DMEM (Welgene) supplemented with 10% inactivated FBS (Welgene), 1% (v/v) penicillin and streptomycin (Lonza) at 37 °C in 5% CO_2._ Transfection of siRNAs was performed using Lipofectamine 2000 (Invitrogen) at a final concentration of 40 nM. siRNAs were obtained from Bioneer with the following sequences: siCon (GCAUUCACUUGGAUAGUAA); NEDD8 (GGAGAUUGAGAUUGACAUU); UBE2M (GAGCUGAACCUGCCCAAGA). All DNA plasmids were transfected using either PEI (Polysciences) or FUGENE 6 (Promega, Madison, WI) according to the manufacturer’s protocol. Arsenite (Sigma, S7400); Clotrimazole (Sigma, C6091); Thapsigargin (Sigma, T9033); Cycloheximide (Sigma, C7698); MLN4924 (Sigma, 5054770001).

### Cloning

eIF3g WT and mutants were cloned into the vector pCI-Neo-FLAG vector using the following primers: eIF3g WT (*Forward primer-GATCCTCGAGATGCCTACTGGAGACTTTGA, Reverse primer-GATCGCGGCCGCTTAGTTGGTGGACGGCTT*); eIF3g ΔRRM (*Forward primer-GATCCTCGAGATGCCTACTGGAGACTTTGA, Reverse primer-TATAGCGGCCGCTTAGTTGTCGTCGGCTCTG*); eIF3g RRM only (*Forward primer-TATCCTCGAGATGGCCACCATCCGTGTCAC, Reverse primer-GATCGCGGCCGCTTAGTTGGTGGACGGCTT*). All other constructs were described previously^24^.

### Western Blotting

Cells were lysed in RIPA buffer (50 mM Tris-Cl (pH 8.0), 150 mM NaCl, 0.1% SDS, 1% NP-40, 1 mM EDTA, 1% Sodium deoxycholate, containing proteinase inhibitors 5 mM NaF, 1 mM PMSF) for 15 min on ice and centrifuged at 14,000 rpm for 12 min. Proteins were quantified using Bradford reagent. Total proteins (20–50 μg) were subjected to SDS–PAGE, transferred to nitrocellulose membranes, and detected with the indicated antibodies. Western blot was performed using an ECL detection system. Antibodies used in this study are listed in **Supplementary Table S2**.

### Immunofluorescence

Cells grown on coverslips were mock-treated or treated with indicated drugs, rinsed twice with PBS (pH 7.4), fixed with 4% paraformaldehyde for 15 min, permeabilized with cold methanol for 10 min, and then blocked in 5% normal horse serum in PBS containing 0.02% sodium azide for 1 h. Cells were then incubated with primary antibodies diluted in blocking solution at room temperature (RT) for 1 h or overnight at 4 °C. After incubation, cells were washed with PBS (three times, 10 min each) and incubated with the appropriate secondary antibodies (Jackson ImmunoResearch ML grade) for 1 h at RT. After incubation, samples were washed three times with PBS (10 min each) and mounted in polyvinyl medium. All images were taken using a Nikon Eclipse 80i fluorescence microscope and processed in ImageJ. At least 100 cells from different fields were counted per condition for quantification.

### Immunoprecipitation

Cells were harvested and lysed in IP buffer (50 mM Tris-Cl (pH 7.5), 150 mM NaCl, 1 mM EDTA, 1% Triton X-100) supplemented with proteinase inhibitors 1 mM PMSF, 10 μg ml^−1^ aprotinin, 5 μg ml^−1^ leupeptin, 0.5 μg ml^−1^ pepstatin and 5 mM NaF on ice for 20 min, centrifuged at 14,000 rpm for 15 min, and the supernatants were collected in a fresh tube. For immunoprecipitation, 1–2 mg lysate was incubated with 20–30 μl Flag agarose beads overnight at 4 °C. The resulting immunoprecipitates were washed at least three times in IP buffer before boiling with SDS sample buffer. The resulting eluates were blotted against the indicated antibodies.

### NEDDylation assay

*In vivo* NEDDylation was performed as previously described^24^. Briefly, HEK293T cells co-transfected with indicated plasmids for 36–40 h were mock-treated or treated with 0.5 mM arsenite for indicated time points. After treatment, cells were washed twice with 1× PBS and collected in 1 mL PBS. About 10% of the cell suspension was centrifuged, and the cell pellet was lysed in RIPA buffer for Western blot analysis, which served as input. The remaining cell suspension was directly lysed in 6 mL guanidinium buffer (6 M guanidinium-HCl, 0.1 M Na_2_HPO_4_/NaH_2_PO_4_, 0.01 M Tris-HCl, pH 8.0) containing 5 mM imidazole, 0.1% Triton X-100, and 10 mM β-mercaptoethanol for 20 min. About 50 μl of Ni-NTA agarose beads were then added directly to the lysates and incubated for 4 h at RT. After incubation, beads were washed once with 800 μl of guanidinium buffer containing 5 mM imidazole, 0.1% Triton X-100 and 10 mM β-mercaptoethanol, once with 800 μl of urea buffer A (8 M Urea, 0.1 M Na_2_HPO_4_/NaH_2_PO_4_, 0.01 M Tris-HCl pH 8.0) containing 5 mM imidazole, 0.1% Triton X-100 and 10 mM β-mercaptoethanol and three times with 900 μl of urea buffer B (8 M Urea, 0.1 M Na_2_HPO_4_/NaH_2_PO_4_, 0.01 M Tris-HCl pH 6.3) containing 5 mM imidazole, 0.1% Triton X-100 and 10 mM β-mercaptoethanol. His-tagged proteins were eluted by incubating the beads in 50 μl elution buffer (5% SDS, 200 mM imidazole, 0.15 M Tris-Cl pH 6.7, 30% glycerol, 0.72 M β-mercaptoethanol, 0.01% Bromophenol Blue) for 20 min. The eluted proteins were directly resolved by SDS–PAGE and blotted to reveal NEDD8 conjugates.

### Cycloheximide chase (CHX) assay

Cells were treated with cycloheximide at a final concentration of 50 µg/ml for the indicated time points. Following incubation, cells were lysed in RIPA buffer supplemented with protease inhibitors and subjected to Western blotting as described above.

### Statistical analysis

All experiments were repeated at least three times unless mentioned in the legends. Statistical analyses were performed with a two-tailed, unpaired Student’s *t*-test. A *P* value<0.05 was considered statistically significant.

## Author contributions

T.O. conceived the project. A.K.J. performed most of the experiments, acquired the data, and prepared the figures for the manuscript. R.M. performed biochemical experiments. A.R. performed bioinformatic analysis. A.K.J., R.M., A.R., and T.O. wrote the manuscript.

## Supplementary figures

**Fig S1.**
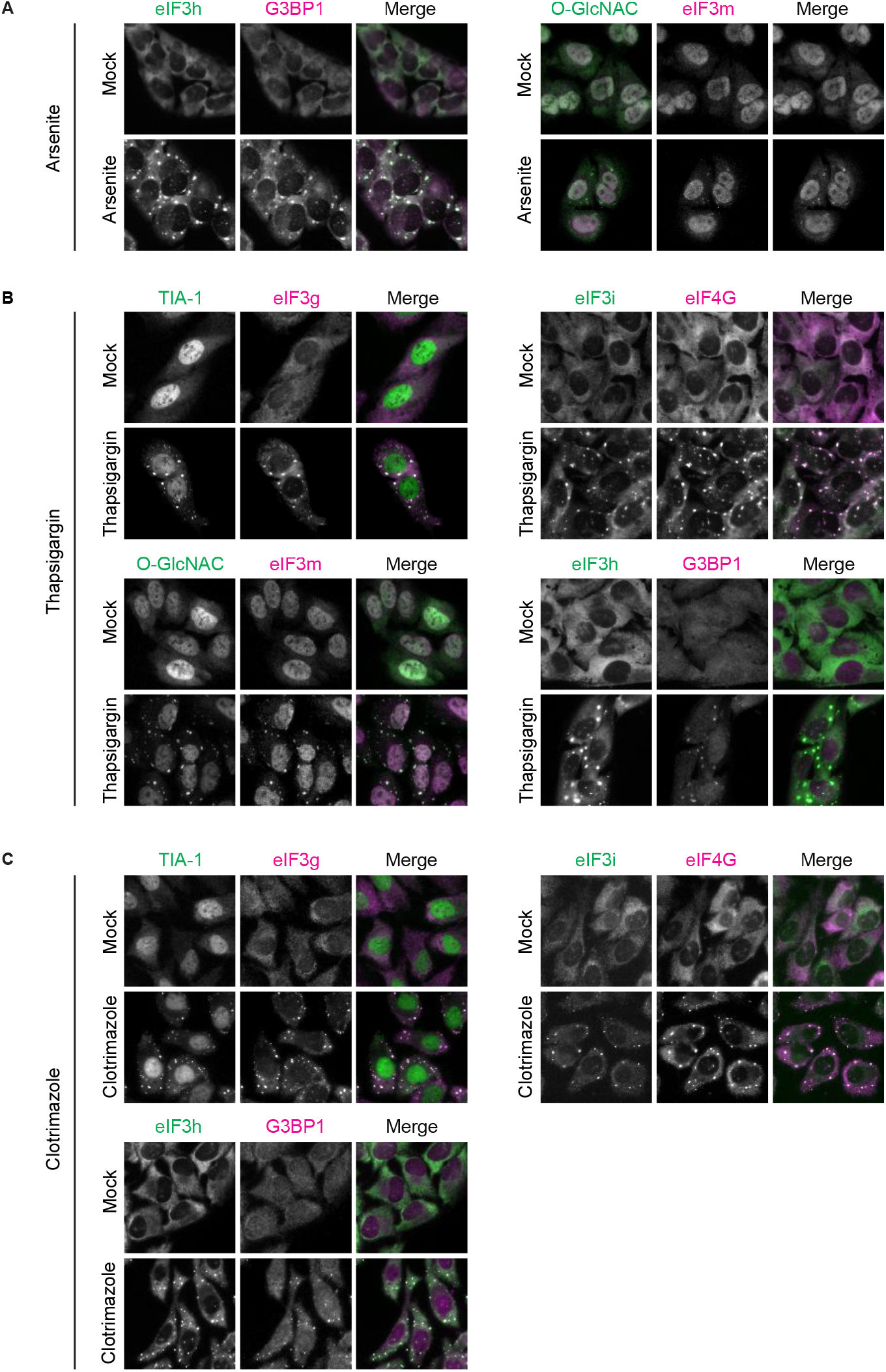
Localization of eIF3g, eIF3i, eIF3m, and eIF3h under different stressors. U2OS cells were treated with (**A**) 0.5 mM arsenite, (B) 1 µM thapsigargin, and (C) 20 µM clotrimazole for 1 h and these cells were fixed and immunostained with the indicated antibodies to reveal SGs. Scale bar, 10 µm.

**Fig S2.**
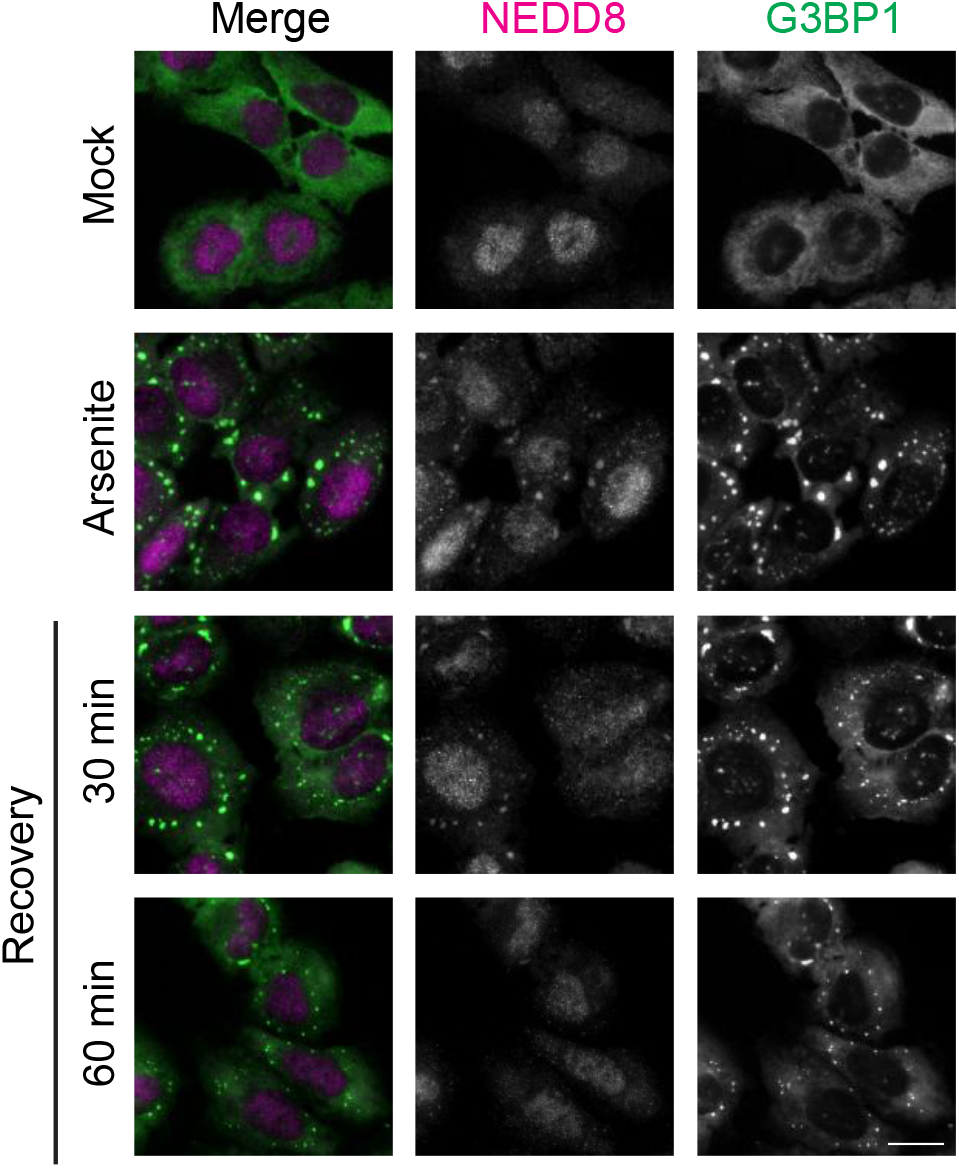
NEDD8 exits SGs early during stress recovery. U2OS cells were treated with 0.5 mM arsenite for 30 min and allowed to recover either 30 min or 60 min, followed by immunostaining with G3BP1 and NEDD8 antibodies. Scale bar, 10 µm.

**Supplementary Table S2.**

| Antibody | Catalog No. | Source | Dilution |  |
| --- | --- | --- | --- | --- |
|  |  |  | WB | IF |
| β-actin (AC-15) | ab6276 | Abcam | 1:5000 | |
| p-eIF2α | BML-SA405 | Enzo | 1:1000 | |
| eIF3g | A301-757A | Bethyl | 1:1000 | 1:500 |
| eIF4G (H-300) | sc-11373 | Santa Cruz |  | 1:500 |
| FLAG | F3165 | Sigma | 1:1000 |  |
| FLAG | F7435 | Sigma |  | 1:500 |
| G3BP1 (H-10) | sc-365338 | Santa Cruz | 1:1000 | 1:500 |
| HA (Y-11) | sc-805 | Santa Cruz | 1:1000 |  |
| NEDD8 | 2745 | Cell signaling | 1:1000 | 1:200 |
| O-GlcNAc | MA1-072 | Thermo |  | 1:1000 |
| His-probe (G-18) | sc-804 | Santa Cruz | 1:500 |  |
| eIF3m (V-21) | sc-133541 | Santa Cruz |  | 1:500 |
| TIA-1 (C-20) | sc-1751 | Santa Cruz |  | 1:500 |
| eIF3i (A-7) | sc-374156 | Santa Cruz | 1:1000 | 1:500 |
| eIF3h | 11310-1-AP | Proteintech |  | 1:500 |
| UBE2M (EPR5333) | ab109507 | Abcam | 1:5000 |  |

